# Machine learning assisted ligand binding energy prediction for *in silico* generated glycosyl hydrolase enzyme combinatorial mutant library

**DOI:** 10.1101/2022.11.29.518414

**Authors:** Igor Guranovic, Mohit Kumar, Chandra K. Bandi, Shishir P. S. Chundawat

**Affiliations:** Department of Biomedical Engineering, Rutgers The State University of New Jersey, Piscataway, New Jersey 08854, USA; Department of Chemical and Biochemical Engineering, Rutgers The State University of New Jersey, Piscataway, New Jersey 08854, USA

**Keywords:** Autodock Vina, In-silico mutagenesis, Multiple Sequence Alignment, Carbohydrate-Active enZyme (CAZy) retriever, Molecular docking simulations, Neural Network, Machine Learning, Glycans, Glycosyl Hydrolase, Glycosynthase

## Abstract

Molecular docking is a computational method used to predict the preferred binding orientation of one molecule to another when bound to each other to form an energetically stable complex. This approach has been widely used for early-stage small-molecule drug design as well as identifying suitable protein-based macromolecule residues for mutagenesis. Estimating binding free energy, based on docking interactions of protein to its ligand based on an appropriate scoring function is often critical for protein mutagenesis studies to improve the activity or alter the specificity of targeted enzymes. However, calculating docking free energy for a large number of protein mutants is computationally challenging and time-consuming. Here, we showcase an end-to-end computational workflow for predicting the binding energy of pNP-Xylose substrate docked within the substrate binding site for a large library of combinatorial mutants of an alpha-L-fucosidase (*Tm*Afc, PDB ID-2ZWY) belonging to *Thermotoga maritima* glycosyl hydrolase (GH) family 29. Briefly, *in silico* combinatorial mutagenesis was performed for the top conserved residues in *Tm*Afc as determined by running multiple sequence alignment against all GH29 family enzyme sequences downloaded from an in-house developed Carbohydrate-Active enZyme (CAZy) database retriever program. The binding energy was calculated through Autodock Vina with pNP-Xylose ligand docking with energy minimized *Tm*Afc mutants, and the data was then used to train a neural network model which was also validated for model predictions using data from Autodock Vina. The current workflow can be adopted for any family of CAZymes to rapidly identify the effect of different mutations within the active site on substrate binding free energy to identify suitable targets for mutagenesis. We anticipate that this workflow could also serve as the starting point for performing more sophisticated and computationally intensive binding free energy calculations to identify targets for mutagenesis and hence optimize use of wet lab resources.

## Introduction

The process of molecular docking simulates noncovalent interactions and predicts binding affinity between two molecules as well as predicts the three-dimensional structure of macromolecular complexes (Morris & Lim-Wilby, 2008). The docking process typically involves three steps: (1) a search for the binding site on the receptor molecule (i.e., protein or enzyme target), and (2) a search for the optimal binding conformation between ligand and target, (3) calculation of binding free energy based on appropriate scoring functions (A. N. Jain & Nicholls, 2008). Such molecular interactions are guided by the molecular shape as well as hydrophobic, electrostatic, van der Waals interactions along with hydrogen bonding, and sum of all termed docking score represents the potentiality of non-covalent binding forces at play (Alberg & Schreiber, 1993). This information can be used to design new drugs that binds to protein targets and also to study binding affinities of ligands to other types of biomolecules, such as RNA and DNA. Hence docking simulations are widely used in the pharmaceutical drug design industry in virtual screening for potential hit identifications from chemical databases, lead optimizations, structure–activity studies, and drug-DNA interactions in recent times (Ferreira et al., 2015; Gschwend et al., 1996; Shoichet et al., 2002). With six degrees of translational and rotational freedom for both ligands and proteins, multiple algorithms have been developed to study their dynamic binding interplay (Huang et al., 2006; Pagadala et al., 2017; Thomsen & Christensen, 2006). Among these workflows, Autodock Vina is a widely used software that uses sophisticated gradient optimization algorithms and has significantly better accuracy of binding mode prediction (Trott & Olson, 2009). Moreover, Autodock Vina has been routinely used in glycoscience field particularly to identify substrate binding regions in the case of glycosyl hydrolases (GHs) and Glycosyltransferases (GTs) (Alsina et al., 2021; Bandi et al., 2021; Pozzo et al., 2014; Xu et al., 2021).

There is significant literature where molecular docking has been used extensively to identify target hits in virtual screening for ligands in drug design (Cross et al., 2009; Gschwend et al., 1996; Hevener et al., 2009; Ma et al., 2011; Onodera et al., 2007; Schneider & Böhm, 2002). On other hand, there are not many studies exploring in-silico generated protein mutants coupled with molecular docking to identify ‘better’ mutants that are amenable to more precise or ‘tighter’ binding (Chiappori et al., 2009). But in broad protein engineering field, structure-based mutagenesis is a common technique to study protein-ligand interactions and protein characteristics under the realm of rational engineering strategies. Several mutagenesis strategies are employed to study site specific interactions for ligand binding such as alanine scanning mutagenesis (Morrison & Weiss, 2001), site saturation mutagenesis where all possible amino acids mutations are investigated in place of wild type residue (Hulme et al., 2007; Williams et al., 1995). Such mutagenesis techniques have been proved to be beneficial to modulate protein function along with substrate specificity as well as to identify key residues involved in the catalytic mechanisms (Geddie & Matsumura, 2004; Yep et al., 2008). The more logical direction of research from here would be random mutagenesis and study the protein-ligand interactions based on those mutants, the literature has shown how this approach has benefited the many case studies cited here including nuclear receptors (Ćelić et al., n.d.; Lim & Huang, 2007; Smith et al., 2004). A combinatorial random mutagenesis approach is a comprehensive mutagenesis technique where all 20 amino acids are replaced at multiple chosen sites (n) simultaneously giving rise to potentially an astronomically large number of possible protein mutants (20^n^) (Yang et al., 2018). The sheer number of possible variants makes the experimental process arduous, time consuming, and even impossible if number of chosen sites are high (Chiappori et al., 2009). *In silico* mutagenesis and screening may constitute as a potential method to circumvent this problem. By leveraging the capabilities of computing power, it is fairly straightforward to create and test large number of combinatorial mutants in order to limit the number of mutants to be tested experimentally. Such computational protein screening strategies have been successfully used in protein engineering for various applications, to name a few; increase enzyme activity, develop bacterial resistance towards antibiotics, improve specificity and engineer novel protein functions (Dahiyat & Mayo, 1996; Hayes et al., 2002; Tokunaga et al., 2008). The novel mutants with intended property identified during *in silico* screening can be synthesized into plasmids and transformed into bacteria to perform activity assays with the purified proteins of those mutants. However, performing Autodock with such high number of combinatorial mutants is also a challenge even with reasonable computing power.

Here, we have utilized a neural network model to predict binding energy based on training data generated from docking of *Tm*Afc mutants with pNP-Xylose. Currently, to the best of our knowledge, there are not any studies in CAZyme field where protein mutants are rapidly screened using a high throughput docking strategy. To give brief descriptions about glycans, they are the most abundant organic molecules in nature, are simple and complex carbohydrates found in free form as well as attached to non-glycan moiety such as proteins, lipids, RNA. They are known to play major structural, metabolic, and physical roles in all cellular systems as they appear to be ubiquitous in all life forms, consequently their applications range from nutrition, regulatory roles, cell to cell interaction, host pathogen interaction to diagnostic and therapeutic roles (Adamczyk et al., 2012; Dube & Bertozzi, 2005; Flynn et al., 2021; Hudak & Bertozzi, 2014; Lauc et al., 2014; Varki & Gagneux, 2015). Every domain of life relies on glycans to mediate a number of biological processes. In order to study these glycans and enzymes associated with them, scientists have created the CAZy database (www.cazy.org) for all enzymes involved in the synthesis, decomposition, and/or alteration of glycans in nature. All CAZymes are properly curated to different groups based on their broad function such as Glycosyl Hydrolase (GH), Glycosyl Transferase (GT), Polysaccharide Lyase (PL), Carbohydrate Esterase (CE), and other auxiliary activity (AA) (Lombard et al., 2014). As of 30 June 2022, 173 GH, 115 GT, 42 PL, 20 CE and 17 AA groups have been identified. Further classification of these groups into families is based on experimentally characterized proteins and sequence similarity from public databases (Cantarel et al., 2009; Levasseur et al., 2013). Since downloading all sequences associated with a particular enzyme family is time consuming, we further developed our in-house tool that can readily download all CAZyme sequences associated with a particular GH or any specific CAZyme family.

Before performing *in silico* mutagenesis, we downloaded all the sequences for GH29 family from CAZy database with the help of our in-house developed CAZy retriever that is now available on the GitHub repository (https://github.com/IgorGuranovic/sequence_retriever). The sequences associated with unique GenBank numbers retrieved from the NCBI database (Federhen, 2012) were fed to multiple sequence alignment using MAAFT algorithm (Katoh, 2002) to identify the highest conserved residues using Jalview (Waterhouse et al., 2009). The MAAFT, one of most popular techniques for protein sequence alignment, uses a heuristic algorithm to calculate pairwise alignment between all of the sequences in the dataset to generate guide tree and sequences are aligned using progressive alignment method (Katoh & Toh, 2008). A total of six chosen sites were then mutated in Pymol (Yuan et al., 2016) randomly to create 20,000 unbiased mutants to perform Autodock simulations with pNP-Xylose as ligand, while ensuring that docking happens only in the designated grid with the correct orientation. We also did energy minimization of all proteins using Rosetta, a well-known macromolecular modeling software suite widely used for protein structure, design, and docking (Das & Baker, 2008; Rohl et al., 2004). The data generated from these docking simulations was segregated for both training and validation purposes. A neural network model with specified parameters was used for training data to achieve the ability to accurately predict ligand docking binding energy for novel mutants. The neural network model, a widely used model in machine learning is made up of series of interconnected nodes, or neurons, that each have weighted input and output; weights are assigned randomly at first and then updated as the model is trained (A. K. Jain et al., 1996). The model learns by adjusting the weights so that the output of model is closer to correct output for training data (Kriegeskorte & Golan, 2019). We used the open source software library, TensorFlow which uses data flow graphs where nodes represent mathematical operations while graph edges represent multidimensional data arrays (tensors) that flow between them (Abadi et al., 2016). The predicted binding energy from the program was tested against initial Autodock data for accuracy and validation.

We have chosen here alpha-L-fucosidase from Thermotoga maritima (*Tm*Afc) as our protein of interest and pNP-Xylose as our binding ligand or acceptor sugar substrate. The protein structure used from the PDB database had a PDB ID-2ZWY (Cobucci-Ponzano et al., 2009a) which already had β-fucosyl azide donor sugar already docked in its active site. We have taken this structure as precursor for performing all *in-silico* mutagenesis. The motivation for taking this as starting structure is highlighted in the utility of conducting a glycosynthase reaction (Hidaka et al., 2010; Wada et al., 2008). The binding energy for the acceptor sugar substrate will likely play an important role in the glycosynthase reaction mechanism (Ducros et al., 2003; Pengthaisong & Ketudat Cairns, 2014). The glycosynthase is an engineered glycosyl hydrolase that can potentially synthesize bespoke glycans or oligosaccharides as naturally occurring glycosyltransferases. However, GT are often difficult to express in *E. coli* and not economic viable due to costly substrates used for chemoenzymatic synthesis (Weijers et al., 2008). The different mutagenesis strategies can be employed to identify improved mutants for glycosynthase applications and molecular docking of ligands with mutant enzymes is one such strategy. The fucosylated oligosaccharides synthesized by mutant *Tm*Afc such as Human Milk Oligosaccharides (HMOs) have critical prebiotic applications such supporting infant growth, gut-immune function against infections, as well as cognitive development (Bode, 2012).

Here we present a case study on ligand binding free energy prediction based on neural network model using molecular docking data generated through Autodock Vina with *in-silico* combinatorial mutants and glycosynthase reaction substrates (**Figure 1**). This is the first reported end-to-end computational workflow to predict binding energy from *in-silico* generated mutants as well as the first study involving glycolyl hydrolase or CAZymes. We calculated the accuracy of our prediction by comparing the data from Autodock Vina test data. We also provide here a list of favorable mutations to observe protein-ligand interactions and analyze the contributions of various residues to ligand binding at particular sites. The workflow involves performing a combinatorial mutagenesis of all residues involved in ligand binding, followed by assessment of the impact of specific amino acid substitutions on ligand binding affinity via molecular docking simulations. We expect such computational workflows will be utility in identifying potentially beneficial mutations for various applications in the protein engineering and glycoscience research community in general.

**Figure 1:**
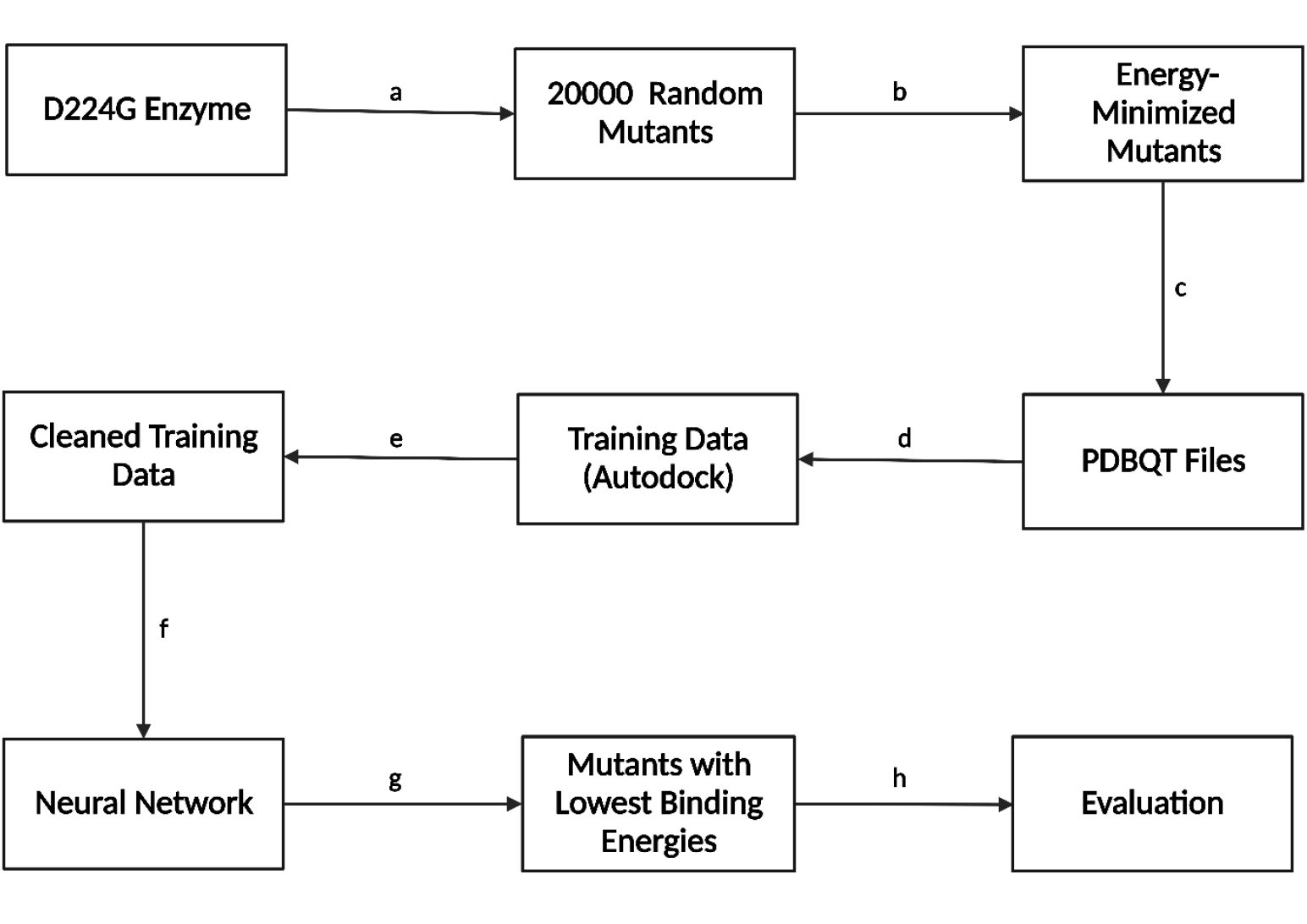
Schematic outlining steps followed in the end-to-end fully automated computational CAZyme-ligand docking workflow starting with *Tm*Afc_D224G as template enzyme sequence, (a) Random mutagenesis of six targeted positions generated using PyMOL, (b) Energy minimization of mutants using PyRosetta, (c) Conversion of PDB into PDBQT files, (d) Autodock simulations of mutant enzymes, (e) Training data cleaning using ligand structural proximity criteria, (f) Training of neural network, (g) Predictions made using neural network, lowest-energy mutants isolated, (h) K-fold cross validation used to evaluate accuracy of neural network model.

## Methods

### CAZy Retriever

This tool was created to retrieve all sequences from a given enzyme class and enzyme family, (for example: Glycoside Hydrolase 29). First, the program identified the GenBank numbers associated with each enzyme in the given class and family, and then used these GenBank numbers to retrieve sequences from NCBI. Each class and family have its own text file on cazy.org encoding these GenBank numbers, and the program parsed this file, storing all GenBank numbers in an Excel file. After that, the GenBank numbers were packaged into a NCBI URL to retrieve FASTA files of the sequences corresponding to the GenBank numbers. The program downloaded FASTA files of 200 sequences at a time. These sequences were packaged into their individual FASTA files and one file of all the sequences.

### Multiple Sequence Alignment

After the sequences from all enzymes of the Glycoside Hydrolase 29 family were packaged into one FASTA file using the CAZy retriever, they were aligned using MAFFT (Katoh et al., 2019). The aligned FASTA file was imported into Jalview, and the consensus value was recorded for each residue in *TmA*fc. The consensus value for a given position in the aligned FASTA file was the percentage of the most frequent amino acid among all enzymes in the file. Next, the amino acid positions of *Tm*Afc with the highest consensus percentage were ranked and displayed as a bar graph (**Figure 5**).

### Autodock Vina

In the active site of *Tm*Afc, there were eight amino acids, namely H34, E66, W67, H128, H129, D224, R254, and E266 that have also been identified previously impacting the glycosynthase reaction (Cobucci-Ponzano et al., 2009b). Since D224G mutation at the catalytic nucleophile site had the highest fucosynthase activity out of all single point mutations of the catalytic nucleophile, D224G was set to be the template enzyme (Agrawal et al., 2021). E266 was not altered since it plays a vital role as acid/base in catalyzing the reaction between the acceptor sugar pNP-xylose and donor sugar β-L-fucosyl-azide (Osanjo et al., 2007). Therefore, the remaining six amino acids were mutated *in silico* in order to identify which permutations of mutations could result in the most effective fucosynthases. Since Autodock takes approximately a minute to complete calculations, this makes it feasible to generate massive amounts of data as opposed to employing molecular dynamics simulations using Amber or LAMMPS (Shirts et al., 2017). However, since there are 20^6^ (64 million) permutations possible, a small subset of these mutants was generated through Autodock and a neural network was trained using on these mutants, with the goal of rapidly screening all 20^6^ mutants.

### In-silico Random Mutagenesis

The six desired amino acids were randomly mutated to generate a mutant PDB file. The template file was the D224G mutant with β-L-fucosyl-azide docked in the active site. The process was repeated 20,000 times, which was carried out in PyMOL using Mutagenesis Wizard. If a duplicate mutant is generated, it is simply deleted from the folder. PyMOL was opened from a Python script, which allowed for automating this process. The heat map in **Figure 2** displays the frequency of a given position being mutated to a certain amino acid, with the vertical axis representing the position and the horizontal axis representing the amino acid. Since the frequencies for all possible mutations are very similar, it can be assumed that the training data set is an unbiased sample of the 20^6^ possible mutants.

**Figure 2:**
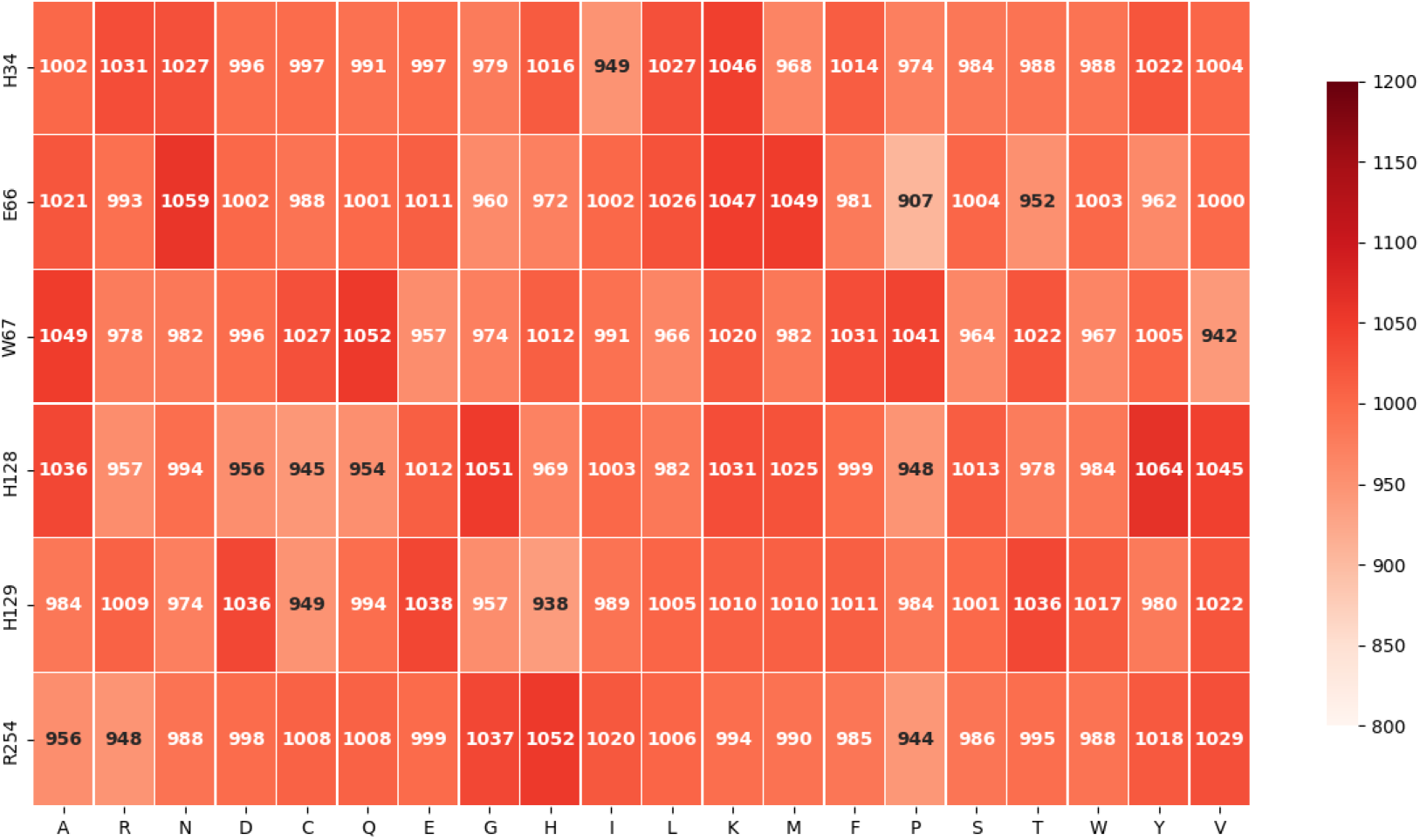
Heat map depicting frequencies of active site positions being mutated to various amino acids for 20000 random mutants; vertical axis depicts position, and horizontal axis depicts mutant amino acid at that specific position.

### Energy Minimization

The mutated structures obtained in PyMOL are not necessarily of the lowest energy. Therefore, PyRosetta was used to get the most stable rotamers for each mutant. 50 Monte Carlo cycles were used in order to maximize structural accuracy while minimizing total runtime (Chaudhury et al., 2010).

### Autodock Docking Automation

Once mutant structures’ energies were minimized, the PDB files were converted into PDBQT file format, which were run using Autodock Vina with a PDBQT file of pNP-Xylose serving as the ligand. The grid box is centered at the active site and does not extend beyond it, meaning that the ligand cannot dock elsewhere in the protein. This way, the network can be trained more accurately since the binding energy would be an accurate representation of the favorability of the active site mutations. All training mutants were run through Autodock Vina, and the binding energies were stored in a Pickle file (file that encodes a Python variable) as well as their corresponding mutants.

### Training Data Cleaning

Even if the grid box is adequately set up, the lowest energy obtained by Autodock might not be the desired energy if the pNP-xylose is not docked in the correct position or orientation. Because of this, a requirement has been introduced to ensure that the Autodock data is representative of the reaction. For each Autodock trial, the program outputted the nine lowest-energy orientations, ordered from lowest to highest energy. The energy associated with a given mutant is the lowest energy where corresponding position satisfies this criterion: At least one of the three hydroxyl oxygens in the pNP-xylose must be no more than 4 angstroms away from C1 carbon in the β-L-fucosyl-azide and no more than 6 angstroms away from both oxygens of the carboxyl group of E266. If neither of the nine positions satisfy this criterion, the Autodock trial is not used as training data. Because of this, there were significantly less than 20000 examples of training data used finally.

### Neural Network

A neural network was created using TensorFlow (Python library) to use this training data generated by Autodock to predict binding energies much faster than using Autodock alone, in a matter of fractions of a second as opposed to approximately a minute. This network has an architecture of four layers in total: one input layer, two hidden layers, and an output layer (**Figure 3**). The 20 possible amino acids for each of the 6 sites were one-hot encoded into 120 input neurons. For an example of a mutant protein, the type of amino acid at a given site was denoted by a 1, while the rest of the input neurons were assigned the value of zero. This is because the input data is categorical, not numerical. The hidden layers contain 64 and 16 neurons respectively, and the ReLU activation function is employed for each hidden layer. The output layer has one neuron with linear activation, representing the binding energy.

**Figure 3:**
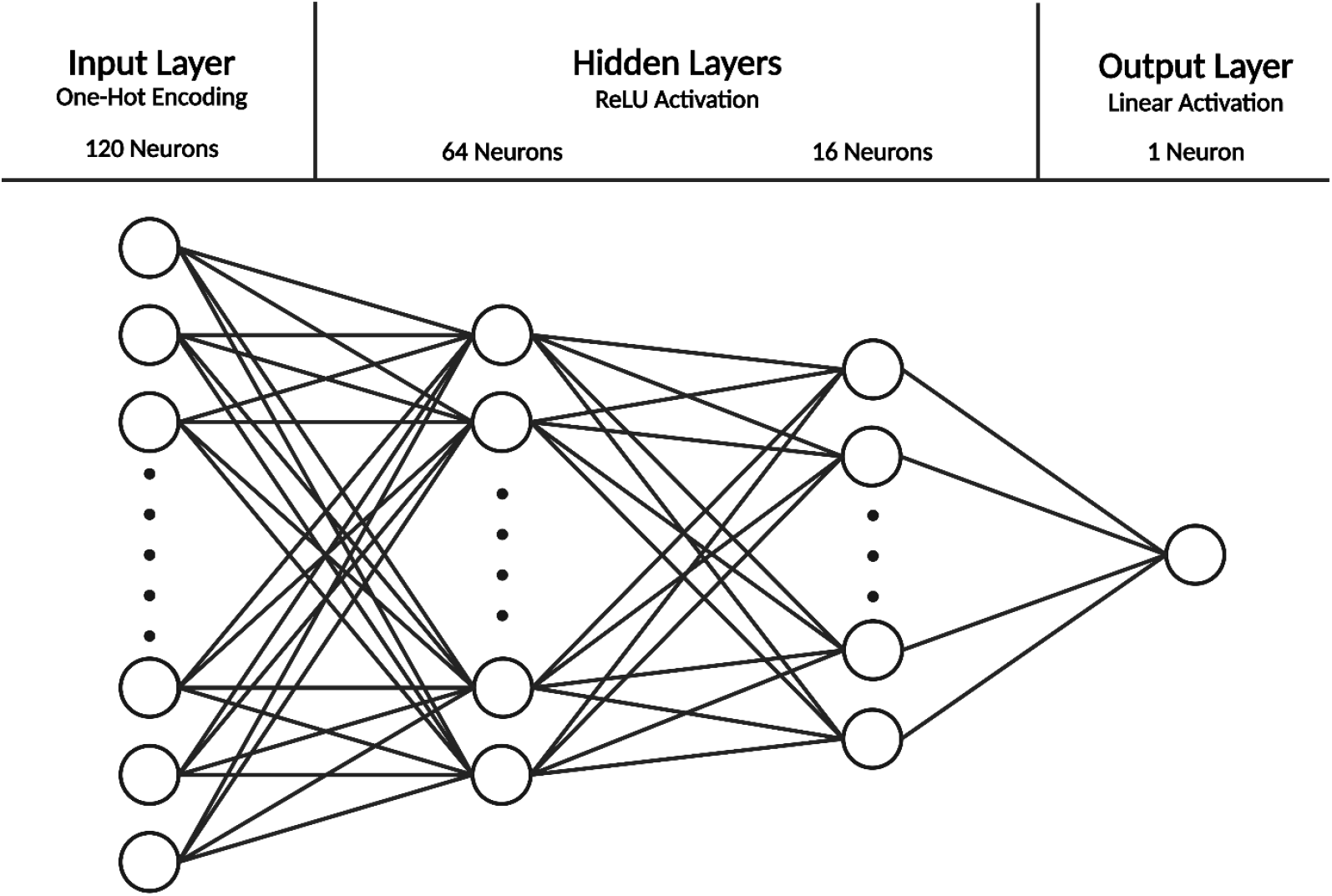
Architecture of the neural network: One-hot encoded input layer with 120 neurons, hidden layers with 64 and 16 neurons respectively (ReLU activation function), and output layer with one neuron encoding the binding energy prediction (linear activation function).

The network used the Adam optimizer, with a learning rate of 0.01, as well as beta1 and beta2 kept at their default values (0.9 and 0.999, respectively). Loss was measured in terms of mean squared error, and 25 epochs were set. Out of the examples of training data, 1/10 were reserved for validation, so only 9/10 were used to train the network.

### Predictions

The neural network model was used to predict binding energies of all possible 20^6^ mutants using the weights and biases obtained from the training data, as well as ranking the mutants in terms of binding energy. In order to reduce the size of the output file, the strategy was to set an arbitrary cutoff energy (in this example 6.2kcal/mol) to filter which mutants were recorded in the file (**Figure 4**). This way, only the most significant mutants were recorded, as opposed to all 20^6^. These mutants can be further tested in the wet laboratory in the form of glycosynthase assays.

**Figure 4:**
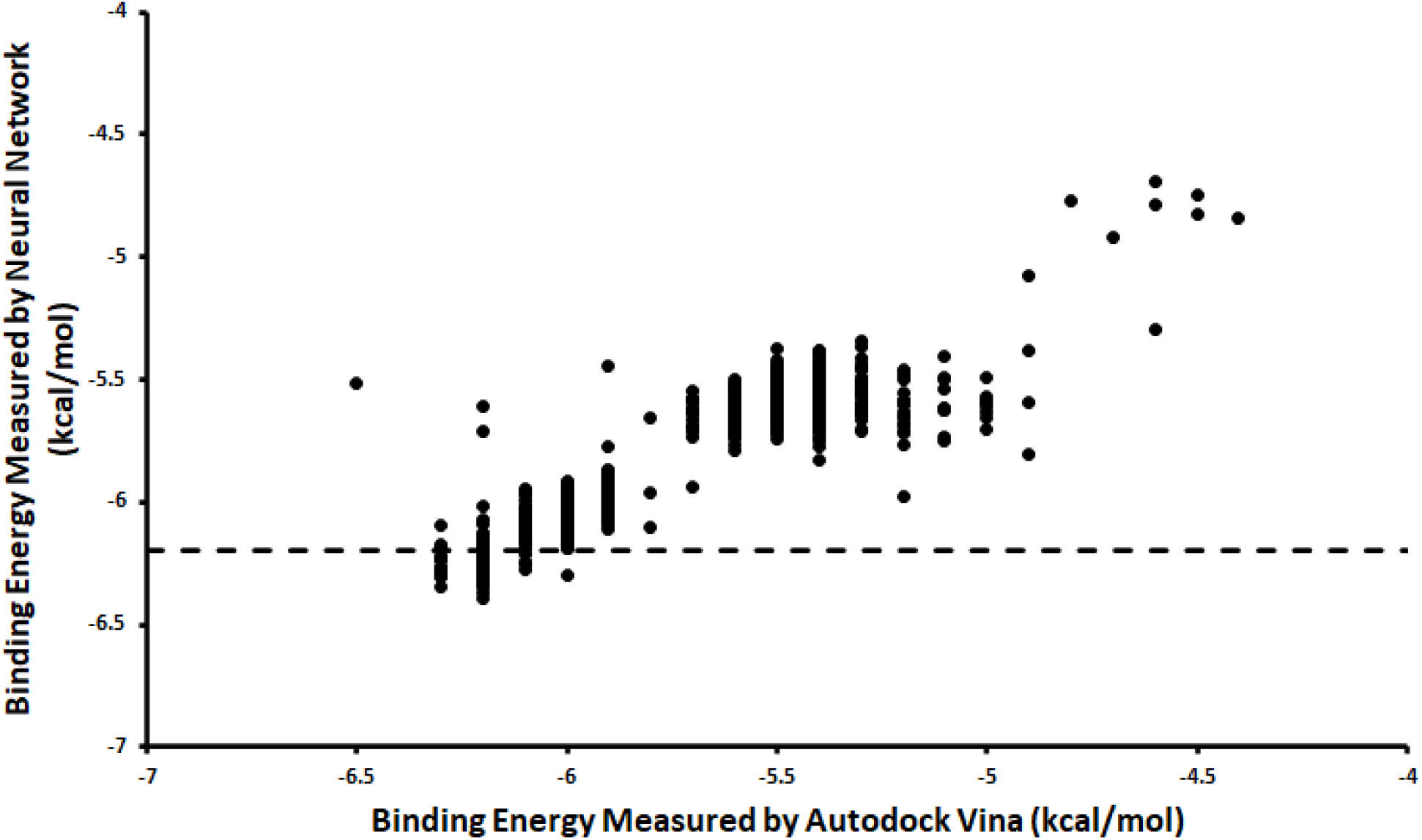
Visualization of accuracy of neural network by testing validation data and comparing predicted energy to the actual one obtained by Autodock. Dotted line represents cutoff energy; only enzymes with predicted binding energies below this energy are recorded in the spreadsheet to limit file size.

**Figure 5:**
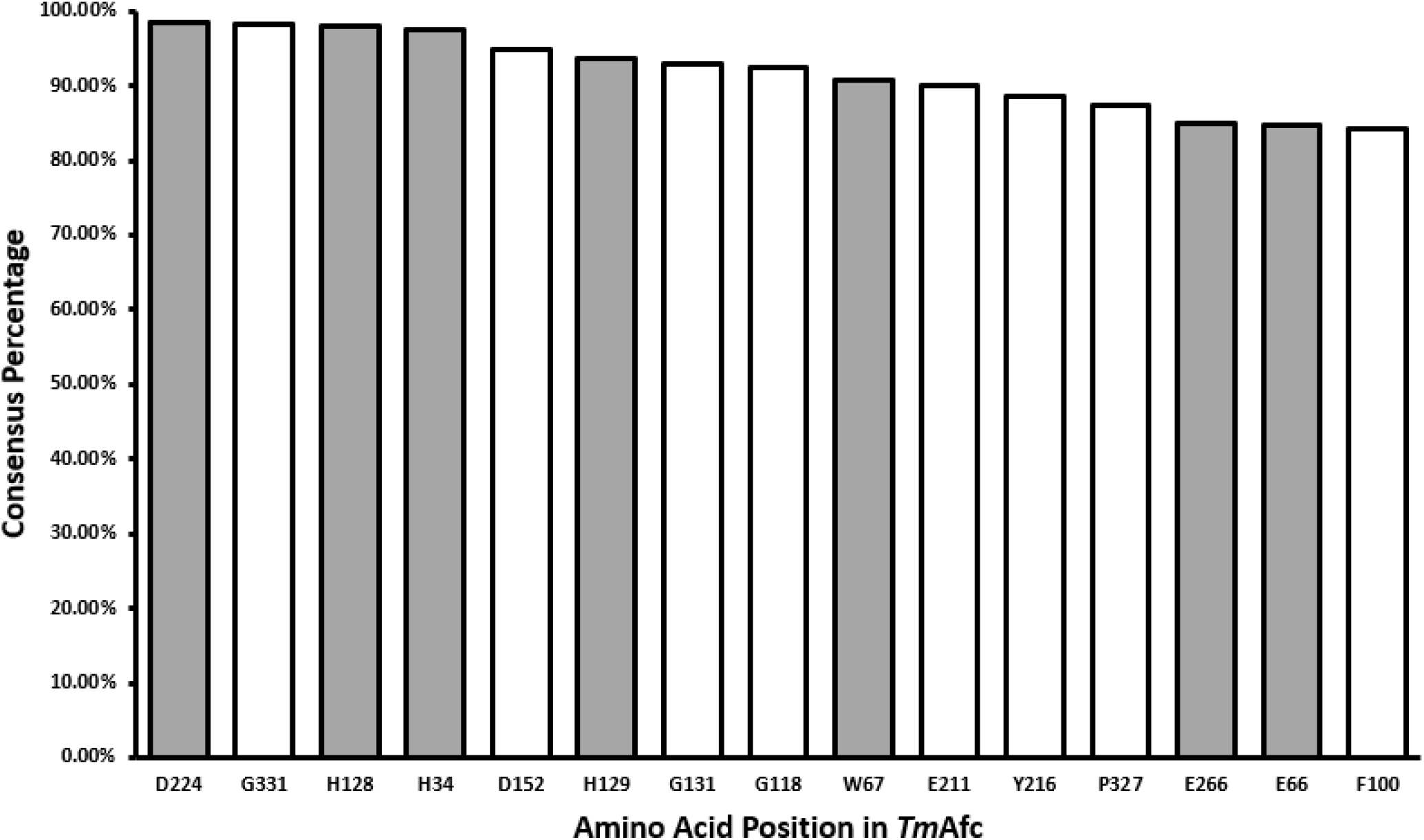
Multiple sequence alignment consensus percentage results for *Tm*Afc against GH29 family enzymes where gray bars represent amino acid positions in the active site of *Tm*Afc.

**Figure 6:**
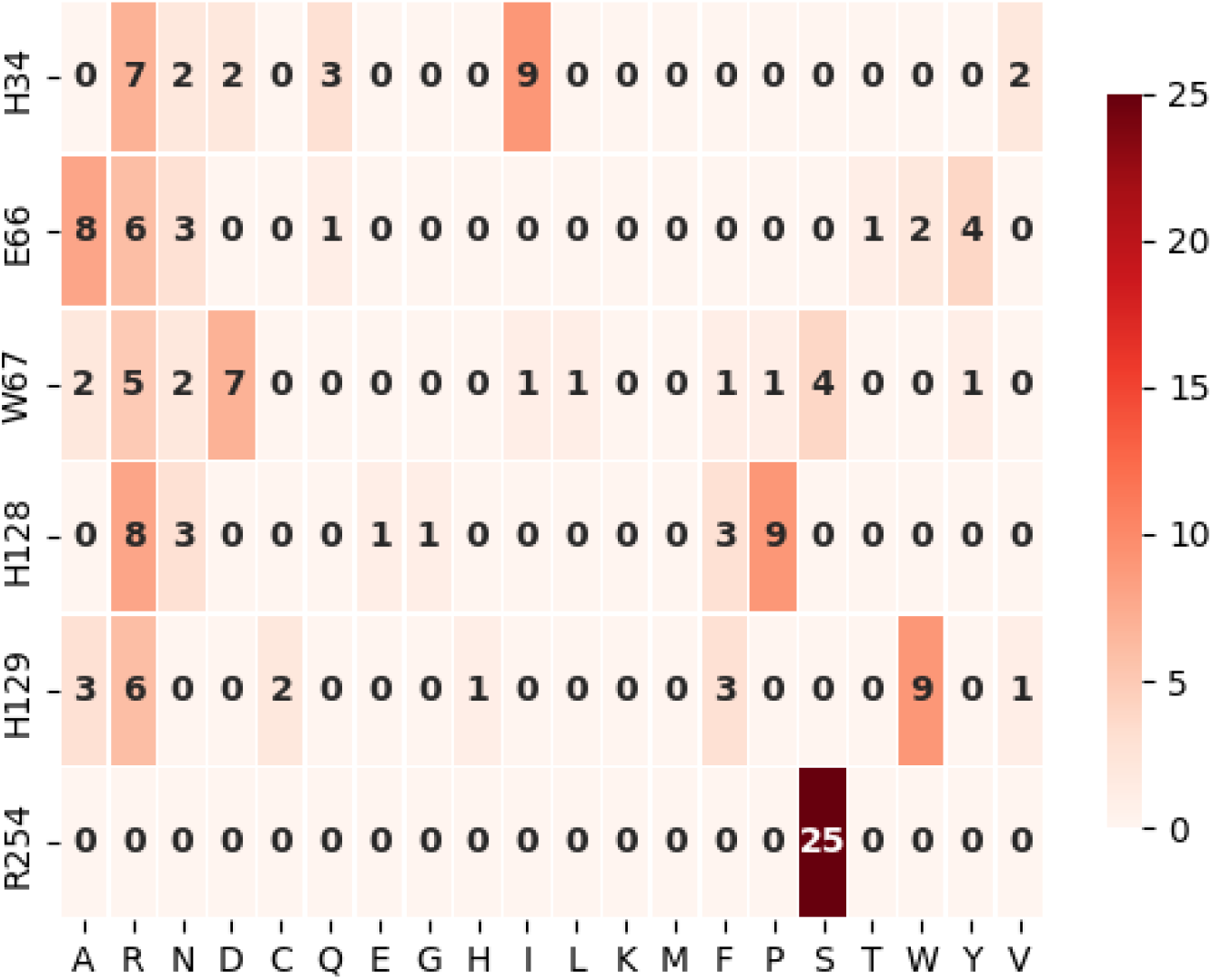
Heatmap showing frequency of mutations among top 25 mutants predicted from neural network model and general trends; H34 tends to favor arginine and isoleucine, E66 tends to favor alanine and arginine, W67 tends to favor aspartic acid, arginine, and serine, H128 tends to favor proline and arginine, H129 tends to favor tryptophan and arginine, R254 favors exclusively serine.

**Figure 7:**
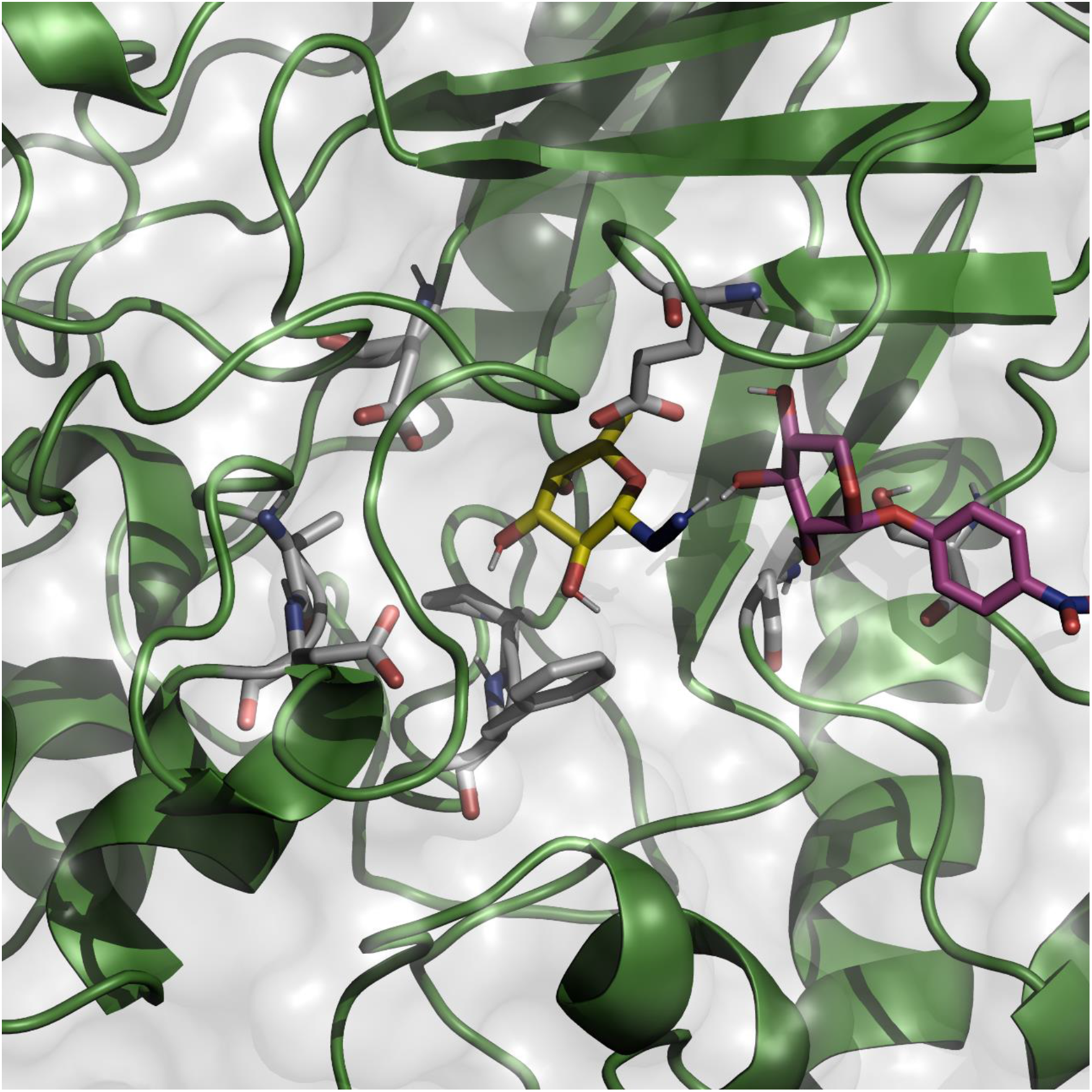
Three-dimensional visualization of docked pNP-xylose (magenta color) and fucosyl azide (yellow color) in the active site (residues shown in light gray) of mutant *Tm*Afc (green color).

## Results

### Multiple Sequence Alignment

Based on the multiple sequence alignment data for *Tm*Afc, seven out of the eight amino acids in the active site were among the 14 highest-consensus amino acids in the entire enzyme, with the exception being R254 as the 90^th^ most conserved amino acid (**Figure 5**).

### Neural Network Evaluation

Once the neural network was trained, the examples of validation data (1/10 of training data sample) were predicted using the network, and the calculated binding energies were compared to the actual energies determined by Autodock. The mean squared error between the two sets of data was 0.02896 kcal^2^/mol^2^, so the root mean squared error is 0.17 kcal/mol. K-fold cross validation was also used to evaluate the network to ensure that the validation data is not biased (**Table 1**). The mean squared error was calculated using 10-fold cross validation, where the data was split into ten folds. Nine of those folds were used for training, one fold was used for validation, and the network was trained ten times such that a different fold was used for validation in each iteration (Emmert-Streib & Dehmer, 2019).

**Table 1:**
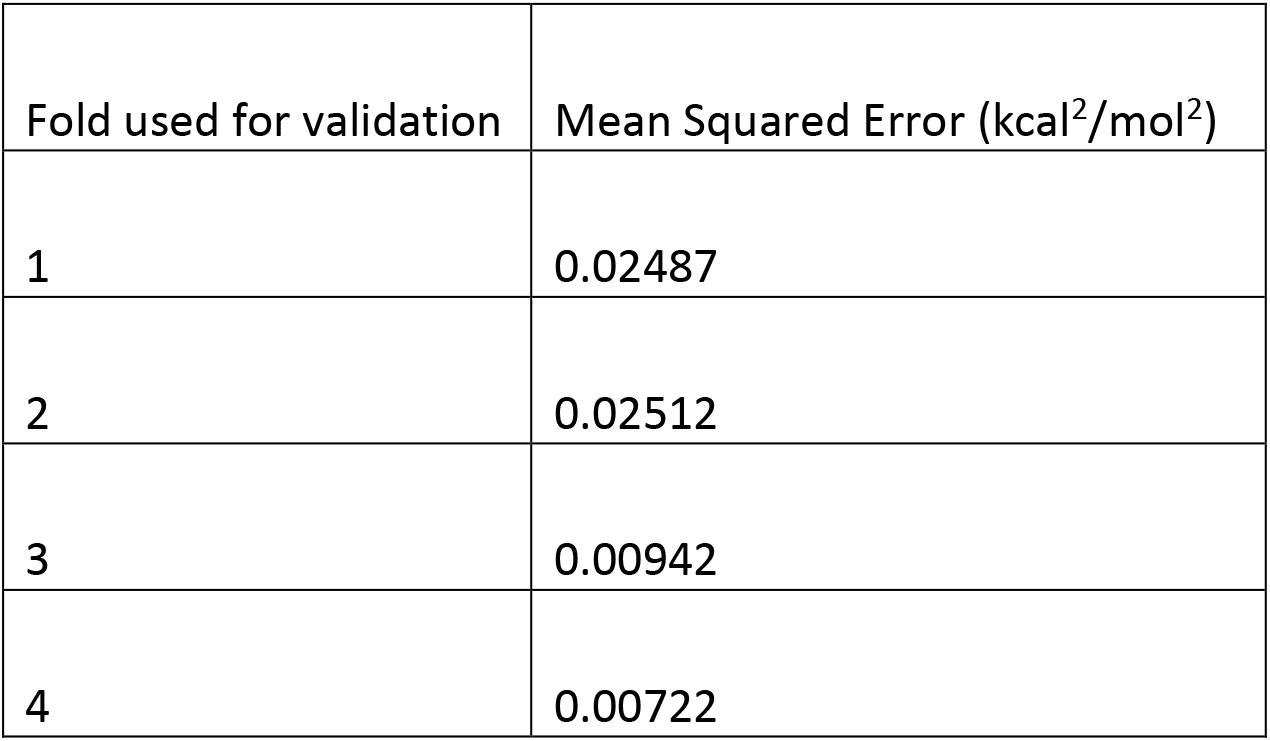
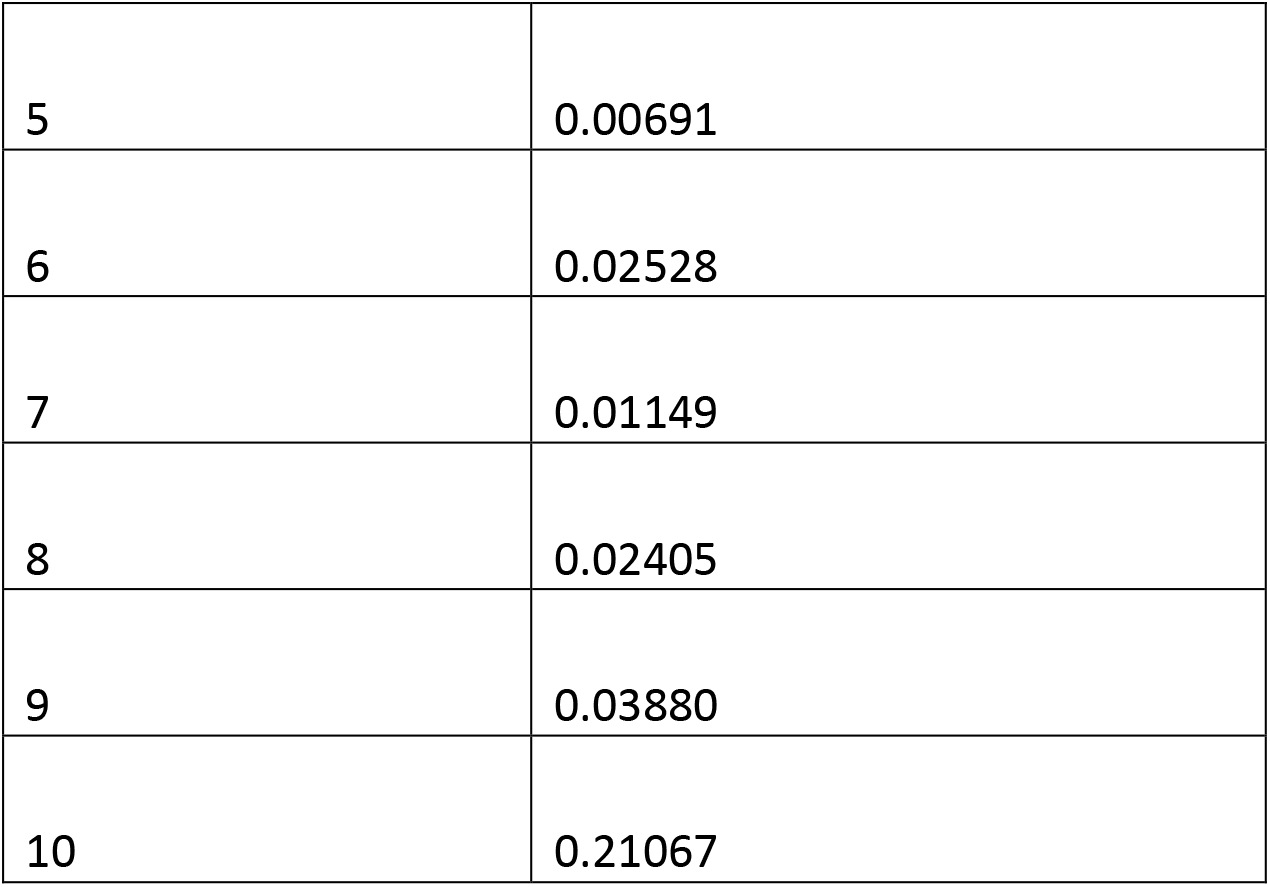
K-fold cross validation of neural network; accuracy for each fold is determined by computing mean squared error (kcal^2^/mol^2^).

### Neural Network Predictions

According to the neural network, when H34, E66, W67, H128, H129, and R254 were free to be mutated, the list of below of 25 mutants were predicted to have the highest binding affinity for pNP-Xylose (**Table 2**). The lowest binding energy was determined to be −6.54386 kcal/mol (Rank 1 mutant).

**Table 2:** Top 25 mutants with lowest binding energies as predicted by the neural network model.

| Rank | Mutant | Predicted Energy (kcal/mol) |
| --- | --- | --- |
| 1 | H34R+E66A+W67D+H128P+H129R+R254S | -6.54386 |
| 2 | H34Q+E66N+W67L+H128N+H129A+R254S | -6.53828 |
| 3 | H34D+E66A+W67D+H128P+H129F+R254S | -6.53367 |
| 4 | H34R+E66A+W67D+H128P+H129F+R254S | -6.53355 |
| 5 | H34I+E66Y+W67R+H128F+H129W+R254S | -6.53061 |
| 6 | H34I+E66R+W67D+H128R+H129W+R254S | -6.53052 |
| 7 | H34D+E66A+W67D+H128P+H129R+R254S | -6.53049 |
| 8 | H34I+E66Y+W67R+H128F+R254S | -6.53034 |
| 9 | H34I+E66R+W67I+H128R+H129W+R254S | -6.52638 |
| 10 | H34Q+E66N+W67Y+H128N+H129A+R254S | -6.52433 |
| 11 | H34R+E66W+W67S+H128E+H129C+R254S | -6.52304 |
| 12 | H34I+E66R+W67N+H128R+H129W+R254S | -6.52279 |
| 13 | H34I+E66R+W67A+H128R+H129W+R254S | -6.51756 |
| 14 | H34N+E66A+W67D+H128P+H129R+R254S | -6.51692 |
| 15 | H34I+E66Y+W67N+H128R+H129W+R254S | -6.5161 |
| 16 | H34R+E66A+W67S+H128P+H129R+R254S | -6.51587 |
| 17 | H34Q+E66N+W67P+H128N+H129A+R254S | -6.51533 |
| 18 | H34R+E66W+W67S+H128G+H129C+R254S | -6.51491 |
| 19 | H34N+E66A+W67D+H128P+H129F+R254S | -6.5142 |
| 20 | H34R+E66Q+W67S+H128P+H129R+R254S | -6.51319 |
| 21 | H34I+E66R+W67R+H128R+H129W+R254S | -6.5117 |
| 22 | H34I+E66Y+W67R+H128F+H129V+R254S | -6.5095 |
| 23 | H34V+E66R+W67A+H128R+H129W+R254S | -6.50943 |
| 24 | H34V+E66T+W67R+H128R+H129W+R254S | -6.50837 |
| 25 | H34R+E66A+W67F+H128P+H129R+R254S | -6.50819 |

## Discussion

We show here how to develop a comprehensive computational pipeline that can predict the binding energy of molecular docking between a large library of protein mutants and ligand substrates using a neural network model. The pipeline is organized into three separate stages for 1) downloading and running MSA for a particular enzyme against its GH family, 2) *in silico* combinatorial mutagenesis and binding energy data generation from Autodock, and finally 3) machine learning model development and checking its accuracy along with analysis of improved outcomes.

These different segments can be altered based on the protein one is interested in working with. *In-silico* mutagenesis does not need knowledge of past information as long as PDB structure and sites are identified based on a rationale to mutate combinatorially. Using a machine learning approach to predict binding energy, especially for a large library of protein mutants can save computational power and time needed for performing protein-ligand docking simulations. Although the model predicts binding energy with reasonable accuracy when we tested against the binding energy data generated from Autodock Vina, there are a few limitations associated with this method. First, an assumption being debated is the correlation between binding energy of an enzyme with its appropriate ligand and the free energy at the transition state (Thyme et al., 2009). Also, Autodock Vina, although widely used for calculating binding energy, is not the absolutely perfect approach to measure binding energy based on docking scores, and that must be further validated through experimental observations. It is important to mention that a neural network was created that uses these already imperfect Autodock results as training data. In spite of having these limitations, this computational toolkit will be useful to have a solid understanding of binding energy distribution among various combinatorial mutants. To test the docking accuracy, we sampled a few improved mutants and their three-dimensional structures, and they were found to be in the expected locations in active sites, giving some credence to this approach.

## Acknowledgements

SPSC acknowledges partial funding support from the US National Science Foundation (Chemistry Award No. 1904890) and Rutgers School of Engineering. IG was supported by Rutgers Aresty program and the Department of Chemical and Biochemical Engineering at Rutgers University.

