## Supplementary Material for "Machine learning assisted ligand binding energy prediction for *in silico* generated glycosyl hydrolase enzyme combinatorial mutant library"

**Python code details:** The python code associated with each section of this paper can be found in GitHub repository using the following link: https://github.com/IgorGuranovic/binding_energy_predictor

**Multiple Sequence Alignment (MSA) visualization:**

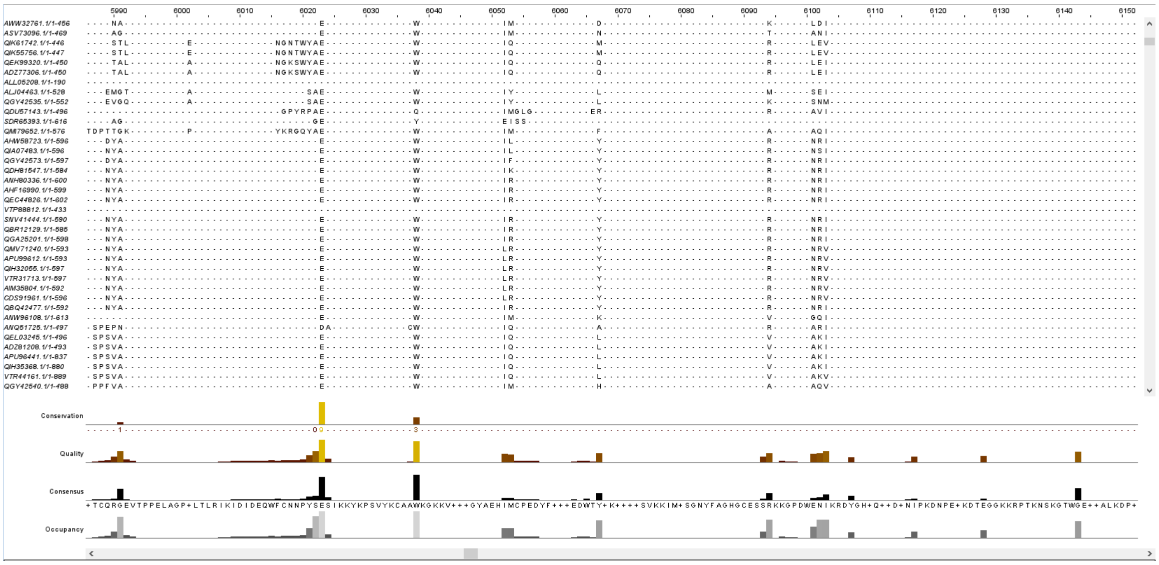


**Figure S1: Multiple Sequence Alignment representational diagram**

**End-to-end simplified procedure to setup and run reported programs:**

Instructions (for Windows 10):

Setup

1) Download PyRosetta folder
2) Install Ubuntu application
3) Download Python 3.7 for Windows, set it to "path" (this will be for PyMOL)
4) Set Python 2.11 (in PyRosetta folder) to "pythonpath" (this will be for AutoDockTools)
5) Download Python 3.8 in Ubuntu (this will be for Rosetta)
6) Pip install all required imports
7) The default wild type enzyme is *Tm*Afc, replace the PDB file with your desired enzyme file and replace all "*Tm*Afc" in the code with the new enzyme file name.
8) Default ligand is pNP-Xylose, replace the PDBQT file with your desired ligand file and replace all "pNP-Xylose" in the code with the new ligand file name.
9) Random Mutagenesis
10) Edit the "aa1.py" file for the desired amount of random mutants for training data (amount pre-set is 20000). The variable is called "num"

11) Open Windows command prompt in the PyRosetta directory and enter “py -3 autodock_automation/aa1.py”
12) The random mutant PDB files are going to be in the folder “PDB_Files”
13) Energy Minimization
14) Edit the “aa2.py” file for the desired amount of Monte Carlo cycles for each mutant, default is 50 cycles.
15) Open Ubuntu command prompt in the PyRosetta directory and enter “python3 aa2.py”
16) The energy minimized mutant PDB files are going to appear in the folder “PDB_Files_Minimized”
17) PDBQT File Preparation + Autodock Trials
18) Open Windows command prompt in the PyRosetta directory and enter “py -3 autodock_automation/aa3.py”
19) Feel free to edit the “config.txt” file to restrict the docking to your desired protein’s active site
20) Training data Pickle files will be in the
21) Neural Network + Predictions
22) Open Windows command prompt in the autodock_automation directory and enter “py -3 autodock_automation/neural_network.py”
23) The network is going to use the training data generated by Autodock
24) A graph will appear in MatPlotLib which shows how accurate the network is
25) Close the graph window if satisfied with network accuracy. If not, modify the architecture, number of epochs, etc. in the “neural_network.py” code and repeat the previous three steps
26) The trained network will predict binding energies for all possible 64 million permutations, and all mutants that are lower in energy than the cutoff (default is -7.6 kcal/mol) ranked in the file “network_predictions.csv”
27) The lowest-energy mutants in this file can be further evaluated through Autodock
28) Validation
29) Open Windows command prompt in the PyRosetta directory and enter “py -3 autodock_automation/aa4.py”
30) Open Ubuntu command prompt in the PyRosetta directory and enter “py -3 aa5.py”
31) Open Windows command prompt in the PyRosetta directory and enter “py -3 autodock_automation/aa6.py”
32) In the “aa4.py” and “aa6.py” files feel free to modify the number (default is 25) of best mutants (ranked by the neural network) to be evaluated by Autodock
33) Once the preceding commands are run, there will be a list of the top 25 mutants in the file “binding_affinity_predictions.csv”, with their corresponding binding energies: one value calculated by the network and the other obtained through Autodock
